# Hippocampal and Midbrain Function in Superagers Relates to Memory for Novelty and Expectation Violation

**DOI:** 10.64898/2026.04.16.718970

**Authors:** Marta García-Huéscar, Linda Zhang, Bryan A. Strange, Darya Frank

## Abstract

Memory performance typically declines with age, but the underlying neurobiological mechanisms remain unclear. Superagers, people over 80 years of age with episodic memory performance comparable to individuals 30 years younger, appear to resist this decline. Novelty and expectation violations are known to engage the hippocampus-midbrain system to enhance memory formation. Here, we examined whether superagers’ superior memory performance is supported by preserved hippocampal-midbrain function during novelty and expectation processing. We manipulated item and contextual novelty (i.e., expectation violations) during encoding to test whether superagers show greater mnemonic benefits than their age-matched peers, whether these benefits reflect enhanced hippocampal and midbrain functioning as measured by fMRI, and whether they are associated with preserved dopaminergic integrity measured with neuromelanin-sensitive MRI. Our results show that, although superagers demonstrated overall superior memory performance, both groups exhibited superior recognition of contextually unexpected items. Nevertheless, differences emerged in the processing of expectation during encoding. Superagers exhibited stronger hippocampal responses to expectation violations and habituation to expected events, irrespective of item novelty. Conversely, typical older adults exhibited reduced midbrain response when expected novelty was absent. Neuromelanin accumulation did not account for group differences in midbrain activity or memory performance. Taken together, these findings suggest superagers benefit from adaptive responses to expectation and its violation, which is therefore a candidate mechanism distinguishing exceptional from typical cognitive ageing.

**SIGNIFICANCE STATEMENT:** Although memory typically declines with age, superagers are individuals aged over 80 who maintain memory performance comparable to people 30 years younger. We examined whether preserved hippocampal-midbrain function during novelty and expectation processing could underlie their superior memory. Superagers exhibited adaptive hippocampal responses to expectation and its violation, with higher activation for unexpected events and habituation to expected events. In contrast, typical older adults showed hippocampal and midbrain responses oriented towards anticipated novel content, despite not showing differences in neuromelanin accumulation. These findings underscore the critical role of hippocampal function in supporting memory preservation in late life and advance our understanding of the neural mechanisms underlying healthy cognitive ageing.

## INTRODUCTION

Memory performance typically declines with age (Glisky, 2007) and is associated with brain atrophy in the medial temporal lobe (Scahill et al., 2003). However, there is mounting evidence that some people can resist this age-related decline. Superagers are people aged over 80 years with the episodic memory performance of individuals 20-30 years younger (Harrison et al., 2012; Rogalski et al., 2013). Superagers have greater grey matter density in the hippocampus (Sun et al., 2016; Harrison et al., 2018; Garo-Pascual et al., 2023), a slower rate of cortical atrophy (Cook et al., 2017; Garo-Pascual et al., 2023) and preserved white matter microstructure (Kim et al., 2020a; Garo-Pascual et al., 2024) compared to age-matched typical older adults. However, how these structural brain differences relate to functional changes and superior memory performance is not well understood. Here, we used task-based fMRI to examine the hippocampus-midbrain system as a key candidate contributing to the preserved memory function in superagers.

Traditionally, ageing research has focused on studying brain structure and resting-state functional connectivity changes over time (Raz, 2000; Damoiseaux, 2017), but these approaches do not directly link brain activity with a specific cognitive function (Mooraj et al., 2025). Task-based neuroimaging can help clarify the cognitive relevance of structural alterations by revealing age-related functional changes (Cabeza et al., 2002; Gazzaley et al., 2005), the effects of neurotransmitter changes in ageing (Berry et al., 2016), and consequently the functional underpinnings of superageing. Indeed, a recent study found preserved neural differentiation in the fusiform and parahippocampal gyri during visual category representations and greater neural reinstatement in superagers (in their 60s) compared to typical older adults (Katsumi et al., 2021).

Novelty engages several neuromodulatory systems, promoting exploratory behaviour and flexible encoding of new experiences (Kafkas and Montaldi, 2018a). One prominent model proposes that the hippocampus detects novelty and signals the midbrain dopaminergic neurons in the substantia nigra and ventral tegmental area (SN/VTA) to release dopamine, promoting plasticity and consolidation of long-term memories (Lisman and Grace, 2005). Contextual novelty, arising from expectation violations (i.e., unexpected pairings of familiar inputs), has been shown to enhance memory performance by engaging hippocampus-midbrain activations during encoding (Kafkas and Montaldi, 2015, 2018b; Frank and Kafkas, 2021). Similarly, item novelty has also been shown to engage midbrain responses and enhance recognition (Bunzeck and Düzel, 2006; Wittmann et al., 2007). Reduced neural responses to familiar items (as repetition suppression) and expectation (expectation suppression) have been associated with predictability, adaptation, and cognitive efficiency (Strange et al., 1999; Yamaguchi et al., 2004; Murty et al., 2013; Vannini et al., 2013; Kim et al., 2020b; Summerfield and de Lange, 2014). By orthogonally manipulating expectation and novelty, we can compare their distinct contributions to adaptive memory formation, i.e., the ability to suppress responses to expected familiar information, while maintaining sensitivity to expectation violations and item novelty.

Furthermore, ageing is associated with impairment in novelty processing (Bunzeck et al., 2007; Schomaker et al., 2022; Steiger et al., 2023) which could be due to age-related deterioration of the dopaminergic midbrain system, leading to reduced dopaminergic synthesis and binding in the brain (Bäckmann et al., 2000; Dahl et al., 2023), thus disrupting reward and expectation violation mnemonic benefits resulting in memory deficits (Düzel et al., 2008). To evaluate this, neuromelanin-sensitive MRI (NM-MRI) is emerging as a modality capable of characterising the functional status of the ageing dopaminergic system *in vivo* (Cassidy et al., 2019; Trujillo et al., 2024).

Understanding the neural and cognitive mechanisms underlying resistance to age-related decline can provide critical insights into potential protective factors against neurodegenerative diseases. Here, we investigated whether superagers’ superior memory performance is driven by a preserved dopaminergic hippocampal-midbrain system compared to typical older adults. By dissociating item novelty from contextual novelty, we aimed to examine their differential and combined contributions to memory formation. We hypothesised that superagers, unlike typical older adults, would benefit from item and contextual novelty (expectation violation), indexed by better memory performance and increased activation in the hippocampus and midbrain. Finally, we explored whether superagers have better dopaminergic function with the use of NM-MRI.

## METHODS

### Participants

Forty-eight healthy older adults from the Vallecas Project in Madrid, Spain (Olazarán et al., 2015) participated in the study. Participants were classified into superager and typical older adult groups based on the same criteria used in previous studies with this cohort (Garo-Pascual et al., 2023, 2024). All participants were right-handed, had normal or corrected-to-normal vision, and no history of any neurological or psychiatric disorders. This project was approved by the Ethics committee of the Universidad Politécnica de Madrid (Reference number 2022-045) and all participants provided written informed consent before taking part in the study. Data from seven participants were discarded: two due to structural brain abnormalities (one superager and one typical older adult), one superager for not meeting the rule-learning criterion, two superagers for non-variable responses in the recognition task (i.e., only pressing ‘new’ or ‘old’), one typical older adult for a high false-alarm rate that led to negative corrected hit rates in both conditions, and one typical older adult due to below chance performance. Final data from 21 superagers (mean age = 82.9, SD = 2.1, 12 females) and 20 typical older adults (mean age = 83.5, SD = 3.1, 13 females) were analysed.

### Stimuli and procedure

Participants performed a memory paradigm consisting of familiarisation, rule-learning, encoding, and recognition phases (Figure 1). Only the encoding phase was performed inside the MRI scanner. The experiment was conducted using PsychoPy version 2021.2.3 (Peirce, 2007).

**Figure 1.**
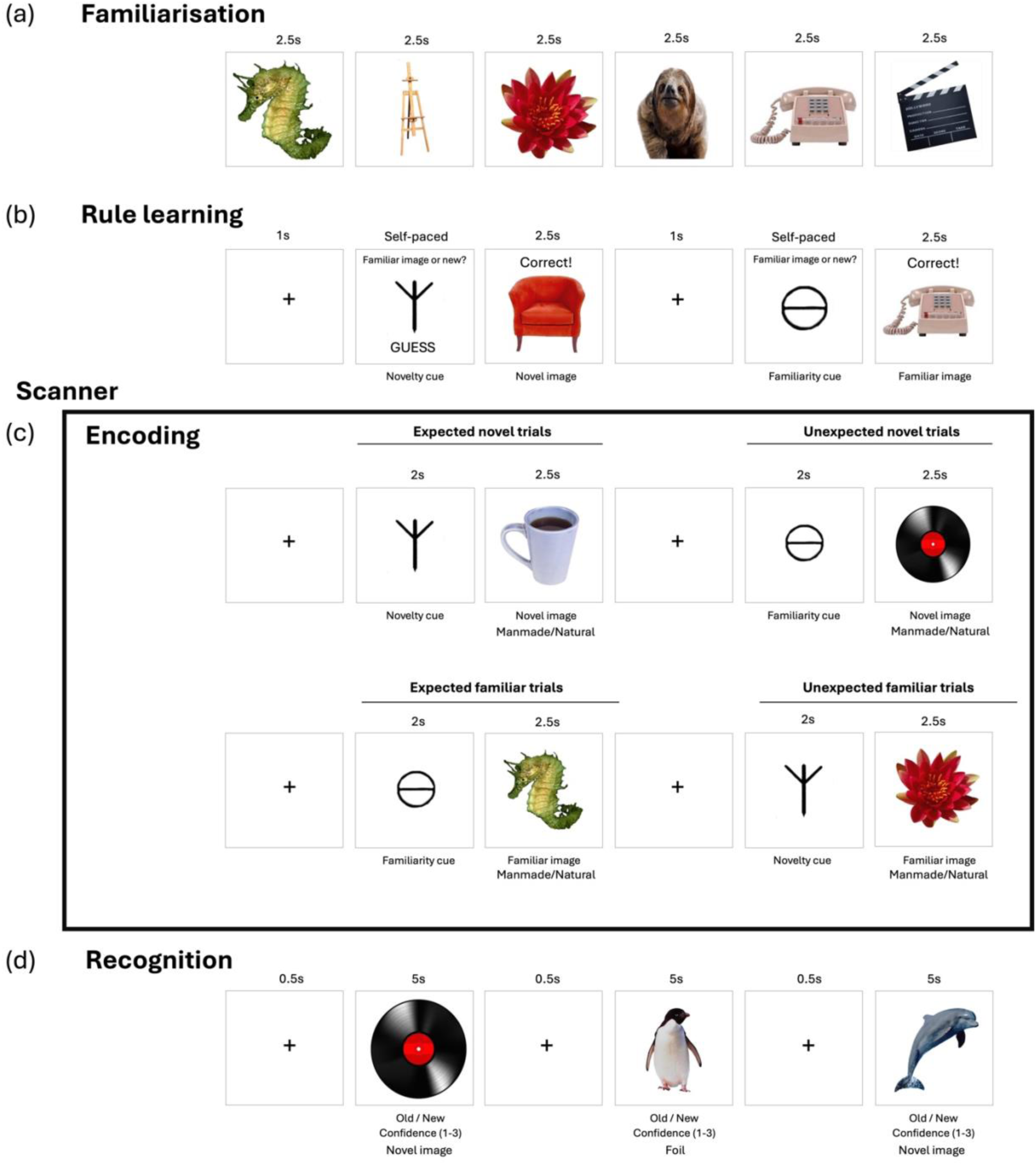
Experimental design. (a) Participants were familiarised with ten images of everyday objects. (b) Participants then had to learn the rule regarding which two familiarity cues preceded a familiar image and which two novelty cues preceded a novel image. (c) Inside the MRI scanner, participants viewed cues followed by images and indicated whether each image depicted a manmade or a natural object. In 30% of the trials, the cue and subsequent image did not match, creating the unexpected familiar and unexpected novel trials. The remaining 70% of trials featured a match between the cue and the image, resulting in the expected familiar and expected novel trials. (d) Participants saw the novel images from the encoding phase mixed with foils and had to indicate if these were old or new, as well as their level of confidence in their response (1 = not sure, 3 = very sure).

#### Familiarisation and rule learning

In the familiarisation phase, participants viewed images of ten objects, each presented five times in pseudo-random order, and were asked to memorise these images to the best of their ability (Figure 1a). Afterwards, they performed a rule-learning task in which they had to learn an association between a cue (an abstract symbol) and an object category – old or new (i.e., whether the image was presented earlier during the familiarisation stage or not). A cue first appeared on the screen and participants were asked to indicate whether they thought the following image would be novel or old. Feedback was then presented on the screen to indicate whether they were correct or not. Participants were instructed that for the first few trials, they would need to guess their response and learn the rule from the feedback they received. There were four cues in total: two familiarity cues predicted the presentation of an old image, and two novelty cues predicted the presentation of a novel image (Figure 1b). Cue symbols were counterbalanced across participants. The rule learning was self-paced (limited to 20 seconds per trial to respond), lasting between 15 to 30 minutes approximately. To ensure effective rule learning, participants were required to achieve at least 75% accuracy in order to proceed to the next phase. Participants who did not reach this criterion repeated the task until the required accuracy was achieved.

#### Encoding

First, participants underwent structural imaging, including a high-resolution T1-weighted MRI and a neuromelanin-sensitive T1-weighted scan (see MRI acquisition section below). Following structural acquisition, participants performed an incidental encoding task consisting of 120 familiar (12 repeats of the 10 familiarised objects) and 120 novel images (not shown previously; Figure 1c) distributed in six experimental runs. Participants were informed that the task would be the same as the rule learning phase, except now they did not need to attend to the cue but were instead asked to indicate whether each image depicted a manmade or a natural object with a button press. The hand participants used to respond with the button box was counterbalanced across participants. Each encoding trial began with a fixation cross jittered between 0.3s and 4.5s, followed by the cue appearing for 2s and the object image appearing for 2s. Critically, to create the unexpected trials, in 30% of the trials there was a mismatch between the cue and the presented image (e.g., a cue that had been paired with novel images was followed by a familiar image). The remaining 70% of trials featuring a match between the cue and the image (e.g., a novelty cue followed by a novel image) are referred to as expected trials. Trials were further categorised as familiar or novel, depending on the type of the outcome image. Participants were presented with six blocks of the encoding task comprising 40 trials each (240 in total), which lasted approximately 30 minutes.

#### Recognition

The recognition phase was performed outside the scanner immediately after the encoding phase (Figure 1d). Stimuli consisted of the novel images seen inside the scanner (120 images) and completely new images to serve as foils (60 images). The ten images used during the familiarisation phase and as ‘familiar’ during encoding were excluded as they were used in both expectation conditions. Participants were presented with a fixation cross for 0.5s and an object image presented for 5s during which they had to indicate if they had previously seen the image or not (i.e., old or new recognition task). After indicating that they had previously seen an image, participants were asked to rate their confidence using a three-point Likert scale (1 = not sure, 3 = very sure). During the debrief, participants were asked if they had noticed the expectation manipulation during the encoding phase (i.e., noticing the mismatched unexpected events).

### MRI acquisition

Scanning was performed in a 3T MRI scanner (Signa HDxt GE) with a phased array eight-channel head coil. Participants had soft pads placed to minimise head movements. First, T1-weighted images (3D spoiled gradient-recalled echo pulse sequence) were acquired with TR = 9.1 ms, TE = 4.1 ms, field of view (FOV) of 100 mm and a matrix size of 256 × 256 with slice thickness of 1mm, yielding a voxel size of 0.9 ×0.9 ×1 mm^3^. Neuromelanin-sensitive T1-weighted images (2D fast gradient spin-echo sequence) were acquired with TR = 600ms, TE = 13.18ms, FOV of 100 mm and a matrix size of 512 × 512 with slice thickness of 2.5 mm, yielding a voxel size of 0.4 × 0.4 × 2.5 mm^3^. Following the structural scans, functional images were acquired using gradient echo-planar imaging (2D single-echo EPI sequence), with TR = 2500 ms, TE = 35 ms, FOV 100 mm and matrix size 64 × 64 with slice thickness of 3mm, yielding a voxel size of 3.4 ×3.4 ×3 mm^3^. Participants performed six blocks of the encoding task inside the scanner (each block lasting 4.2 minutes). Two participants had five fMRI blocks due to a scanner malfunction.

### MRI data analysis

Functional MRI data were preprocessed and analysed using SPM12 (Statistical Parametric Mapping; Wellcome Centre for Human Neuroimaging, University College London; https://www.fil.ion.ucl.ac.uk/spm). Preprocessing included realignment, reslicing, and slice-timing correction to the middle slice. Each participant’s high-resolution T1-weighted anatomical image was co-registered to the mean functional image. T1 images were segmented using the CAT12 toolbox (version r223, Gaser et al., 2024, https://neuro-jena.github.io/cat). Voxel-based morphometry (VBM) analyses were conducted to examine grey matter volume differences, and Threshold-Free Cluster Enhancement (TFCE) was performed using CAT12 with 5000 permutations, with statistical significance assessed at p < 0.05 FWE-corrected. Functional data were spatially normalised to MNI space using the DARTEL toolbox (Ashburner, 2007) implemented in SPM12 and smoothed with a 6 mm FWHM Gaussian kernel. For each participant, a first-level general linear model (GLM) was used to model individual trials (least squares-all approach), representing the conditions of interest (cue, expected familiar, unexpected familiar, expected novel, and unexpected novel). Additional regressors were included to control for motion and physiological noise using CompCor correction (Behzadi et al., 2007). Regions of interest (ROIs) included the bilateral hippocampus, derived from the Harvard-Oxford anatomical atlas, and the midbrain, using a previously published probabilistic SN/VTA mask thresholded at 25% (Pauli et al., 2018). For the linear-mixed effects model (see Statistical Analysis section below), ROI parameter estimates were extracted from the first-level GLM t-maps to reduce noise and enhance detection sensitivity (Dimsdale-Zucker et al., 2018).

Midbrain volumetric data were derived using the Brainstem Substructures toolbox (Iglesias et al., 2015) implemented in FreeSurfer version 7.4.0. NM-MRI data were processed via a robust, automated voxel-wise pipeline (Wengler et al., 2020). Motion correction, averaging, and T1 co-registration were conducted in SPM12, followed by spatial normalisation to MNI space using Advanced Normalization Tools (ANTs). Neuromelanin signal intensity was quantified using the contrast-to-noise ratio (CNR), calculated at each voxel as: CNR_v_ = { [I_v_ – mode(I_CC_)] / mode(I_CC_) } *100. Where mode (I_CC_) was estimated using a kernel-smoothed histogram of all voxels within the crus cerebri (CC) mask, a white matter region with minimal neuromelanin content. The CC mask was manually traced on a NM-MRI template generated from 40 individuals. This mode-based approach was chosen over the mean or median to improve robustness to edge artifacts and outliers (complete details available in Cassidy et al., 2019).

### Statistical analysis

All statistical tests were performed in R software version 4.3.2 (R Development Core Team, 2008). For the ROI analysis of encoding fMRI data, linear mixed-effects models were computed using the lme4 package (Bates et al., 2015). We used mixed effects modelling to account for variability within and across participants and conditions simultaneously. The within-subjects’ factors were expectation (expected vs. unexpected cue condition) and novelty (familiar vs. novel image), and the between-subject factor was group (superagers vs. typical older adults). Participant intercepts were included as random effects (including random slopes for expectation and novelty in the models yielded similar findings). False discovery rate method was used to correct for multiple comparisons. The slopes of each predictor in the model were assessed using the sim_slopes function from the jtools package (Long, 2022). Between-group volumetric analyses were adjusted by total intracranial volume (TIV). For recognition performance, we used participants’ corrected hit rate as a measure of accuracy, calculated as hit minus false alarm rate. A 2 x 2 mixed ANOVA on recognition data was performed, with the within-subjects factor expectation at encoding (expected vs. unexpected) and the between-subject factor group (superagers vs. typical older adults). Post hoc comparisons of estimated marginal means were conducted using Wald z-tests. Extraction and plotting of the effects reported in the results section below were conducted using the effects package (Fox, 2003) and ggplot2 (Wickham, 2009). The data and code used for the analysis is available here: https://github.com/mghuescar/neema2.

## RESULTS

### Demographics and neuropsychological results

We did not observe significant differences between superagers and typical older adults in age, sex, or years of education (Table 1). By definition, superagers outperformed typical older adults in the neuropsychological selection criteria scores, namely the Free and Cued Selective Reminding Test (FCSRT), Animal Fluency test, Digit Symbol Substitution test and 15-item Boston Naming test. Furthermore, no group differences were observed in genetic biomarkers associated with increased (*APOE* ε4 allele) or decreased (*APOE* ε2 allele) risk of Alzheimer’s disease, consistent with previous findings reported in Garo-Pascual et al. (2023).

**Table 1.** Demographic, neuropsychological and APOE genotype differences between superagers and typical older adults.

|  | **Superagers**<br>(n=21) | **Typical older adults** (n=20) | **Statistic** | **P** | **FDR P** |
| --- | --- | --- | --- | --- | --- |
| **Demographics** |  |  |  |  |  |
| Age in years, mean (SD) | 82.9 (2.1) | 83.5 (3.1) | T = 0.7 | 0.48 | 0.48 |
| Sex |  |  |  |  |  |
| Female, n (%) | 12 (57.1) | 13 (65) | X = 0.03 | 0.84† | 0.85† |
| Male, n (%) | 9 (42.8) | 7 (35) |  |  |  |
| Education in years, mean (SD) | 14.3 (5.6) | 11.4 (5.4) | T = -1.7 | 0.09 | 0.09 |
| **Neuropsychological scores** |  |  |  |  |  |
| Free and Cued Selective Reminding Test free delayed recall score, mean (SD) | 12.8 (1.7) | 7.1 (1.5) | T = -10.9 | < 0.001 | < 0.001 |
| Animal Fluency Test total score, mean (SD) | 21 (3.9) | 15.6 (4.0) | T = -4.3 | < 0.001 | < 0.001 |
| Digit Symbol Substitution Test total score, mean (SD) | 21.4 (7.1) | 17.4 (5.2) | T = -2.0 | 0.04 | 0.04 |
| 15-item Boston Naming Test total score, mean (SD) | 14.4 (1.1) | 11.9 (3.0) | T = -3.6 | < 0.001 | 0.0007 |
| Mini Mental State Examination total score, mean (SD) | 29.3 (0.9) | 28.6 (1.1) | T = -2.0 | 0.05 | 0.05 |
| **APOE alleles** |  |  |  |  |  |
| E2/E3 | 3 (14.3%) | 3 (15%) | - | 0.93‡ | 1‡ |
| E3/E3 | 16 (76.2%) | 14 (70%) | - | 0.93‡ | 1‡ |
| E2/E4 | 0 (0%) | 1 (5%) | - | 0.93‡ | 1‡ |
| E3/E4 | 2 (9.5) | 2 (10%) | - | 0.93‡ | 1‡ |
| Data are n (%) or mean (SD). Comparisons done with t-test. †Comparisons done with $\chi^2$ test.<br>‡Comparisons done with Fisher's test | | | | | |

### Behavioural results

For the recognition phase, a 2 x 2 mixed ANOVA (factors: group and expectation) on corrected hit rate showed a main effect of group (F_(1,39)_ = 9.09, p = 0.005, η² = 0.17), with superagers showing better recognition performance than typical older adults and a main effect of expectation (F_(1,39)_ = 7.67, p = 0.009, η² = 0.01), with both groups having better recognition of unexpected compared to expected objects (Figure 2A). We did not find a significant interaction between group and expectation (F_(1,39)_ = 0.06, p = 0.81, η² < 0.001). These findings indicate that superagers exhibit overall superior recognition performance compared to typical older adults, regardless of expectation condition, and both groups demonstrated an enhanced recognition of unexpected events.

**Figure 2.**
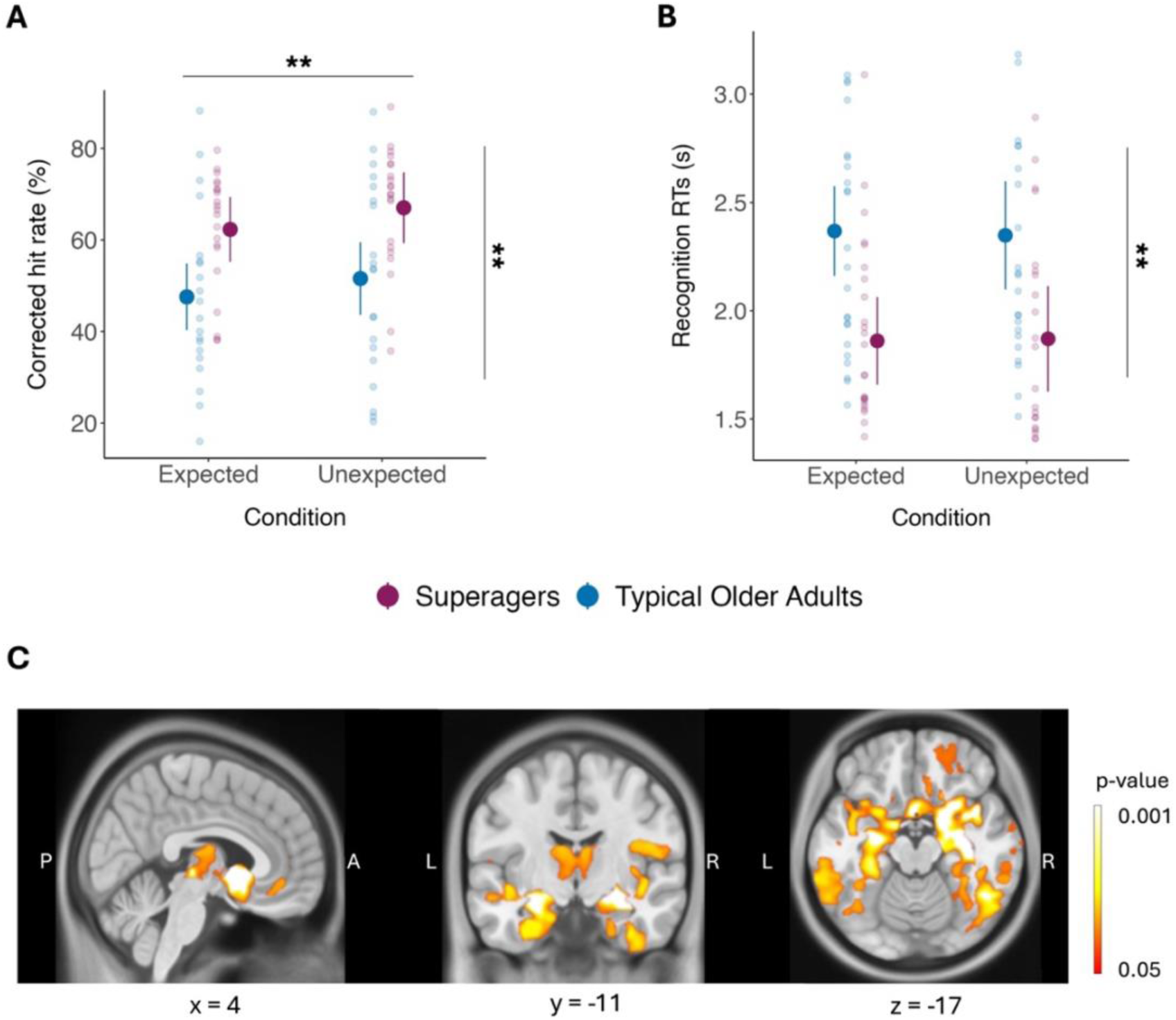
Behavioural results for the recognition phase and neuroanatomical differences between superagers and typical older adults. **A**, Recognition memory performance (corrected hit rate), with superagers remembering more images than typical older adults regardless of their expectation (p = 0.005). Both groups remember more unexpected compared to expected objects (p = 0.009). **B**, Reaction times in seconds (RTs) for the recognition phase, with superagers responding faster than typical older adults in both the expected and unexpected conditions (p = 0.003). **C**, Superagers show larger grey matter volumes in bilateral hippocampus, amygdala, thalamus and nucleus accumbens. TFCE family-wise error-corrected p<0.05. Error bars reflect 95% confidence interval. **p < 0.01

We also analysed the confidence responses for remembered objects with a 2 x 3 mixed ANOVA (factors: group and recognition condition – expected, unexpected or foil) and found a main effect of condition (F_(1,43.79)_ = 24.16, p < 0.001, η² = 0.13), with confidence ratings superior in both groups both for expected and unexpected images compared to foils. We did not find a significant difference between groups (F_(1,39)_ = 1.18, p = 0.28, η² = 0.22) or an interaction between group and condition (F_(1,43.79)_ = 0.37, p = 0.56, η² = 0.002).

For reaction times during recognition, a 2 x 3 mixed ANOVA (factors: group and condition – expected, unexpected or foil) showed a main effect of group (F_(1,39)_ = 9.84, p = 0.003, η² = 0.14), with superagers responding faster than typical older adults (Figure 2B), and a main effect of condition (F_(1,44.47)_ = 12.86, p < 0.001, η² < 0.09), with both groups responding slower to foils compared with expected and unexpected images. We did not find a significant interaction between group and condition (F_(1,44.47)_ = 1.07, p = 0.31, η² = 0.008). There was no significant effect of expectation when foils were excluded from the analysis (F_(1,39)_ = 0.02, p = 0.88, η² < 0.001). These results suggest that, irrespective of encoding expectation, superagers responded faster than typical older adults during recognition.

Performance during the rule learning phase (number of sessions required to learn the rule), and accuracy and reaction times during the encoding phase (manmade/natural decisions) are provided in Supplemental Materials. Superagers required fewer sessions to learn the rule but were slower than typical older adults when making the manmade/natural decision (Table S1).

### Neuroanatomical results

VBM analysis (Figure 2C) revealed that superagers have greater grey matter volume in regions of the medial temporal lobe (bilateral hippocampus, parahippocampal and entorhinal cortices), bilateral thalamus, nucleus accumbens, amygdala and basal ganglia (TFCE FWE-corrected p < 0.05). These findings are consistent with previous observations from the larger cohort of superagers and typical older adults from which the present sample was selected (Garo-Pascual et al., 2023). In contrast, volumetric analysis of the midbrain revealed no significant differences between superagers and typical older adults (t_(39)_ = 0.02, p = 0.87, η² < 0.001).

### fMRI Results

Given our a priori hypothesis, we focused our analysis on the ROIs of interest (hippocampus and SN/VTA). Additionally, we report univariate whole-brain exploratory analysis in the Supplemental Materials, which revealed greater activation in the superior parietal lobe in superagers compared to typical older adults for the main effect of expectation violations (Figure S1).

#### Superagers show hippocampal habituation to expectation

We first examined hippocampal activation to assess the effects of expectation and novelty using a 2 x 2 x 2 linear mixed effects model (factors: group, expectation, and novelty). No significant effects were found for the main effects of group (β = -0.01, χ²(1) = 2.51, p = 0.11), expectation (β = 0.003, χ²(1) = 0.63, p = 0.42) and novelty (β = -0.005, χ²(1) = 1.59, p = 0.20). There was a trend towards an interaction of expectation and novelty (β = -0.007, χ²_(1)_ = 3.40, p = 0.06). The rest of the interactions were not significant (all p’s >0.17). Subsequent tests revealed that both groups had higher activation for expected novel compared to expected familiar events (t = -3.10, p _FDR =_ 0.01), suggestive of a preserved hippocampal processing of item novelty when it is expected. The rest of the contrasts were not significant (all p’s > 0.18).

The hippocampus has been shown to express repetition suppression effects (Kim et al., 2020b), indicative of habituation to repeated presentations, a response which declines with age (Pihlajamäki et al., 2011; Adams et al., 2021). Repetition suppression can be interpreted as an adaptive mechanism through which the brain becomes more efficient at processing previously encountered information. In the context of our task, successful repetition suppression would be manifested in a reduction in hippocampal activity for repeated familiar items as the task progresses (and conversely, failure would be increased activity to repeated items). In addition to the traditional operationalisation of repetition suppression, our design enables further examination of the temporal dynamics of expectation processing, and whether expected outcomes also produce expectation suppression effects (i.e. reduced activity for expected items). To examine this, we added the experimental run (six encoding blocks of 40 trials each) to the linear mixed-effects model (factors: group, expectation, novelty and run). We found a significant main effect of run (β = –0.022, χ²_(5)_ = 12.56, p = 0.028), and a significant interaction between group and run (β = – 0.027, χ²_(5)_ = 20.96, p < 0.001), suggesting that the pattern of hippocampal activity over time differed between groups as a function of time. Trends toward significance were observed for the interaction between expectation and novelty (β = –0.007, χ²_(1)_ = 3.32, p = 0.069) and the three-way interaction between expectation, novelty, and run (β = 0.01, χ²_(5)_ = 9.24, p = 0.09), suggesting a potential modulation of hippocampal responses by expectation and novelty across runs. Importantly, a significant three-way interaction between group, expectation, and run was observed (β = –0.023, χ²_(5)_ = 11.28, p = 0.046), indicating that the influence of expectation on hippocampal activity varied over runs with differences between groups.

To unpack this interaction, we split the data by expectation. For expected events (Figure 3A), there was a main effect of novelty (β = –0.005, χ²_(1)_ = 9.45, p = 0.002), a main effect of run (β = –0.022, χ²_(1)_ = 12.44, p = 0.02) and an interaction between group and run (β = –0.027, χ²_(1)_ = 36.45, p <0.001). No other interactions were significant (all ps >0.30). Post hoc comparisons of hippocampal slopes revealed a significant difference between groups for expected events: while typical older adults exhibited increased hippocampal activation across runs for novel objects (t = 3.97, p < 0.001) and a marginal increase for familiar objects (t = 1.93, p = 0.05), superagers instead showed a decrease in hippocampal activation over time for familiar (t = -2.05, p = 0.04) and novel objects (t = -2.22, p = 0.03). We additionally split the data by group to characterise the temporal dynamics within each group. Notably, only the superagers showed a significant expectation-by-run interaction; these results are reported in the Supplemental Materials.

**Figure 3.**
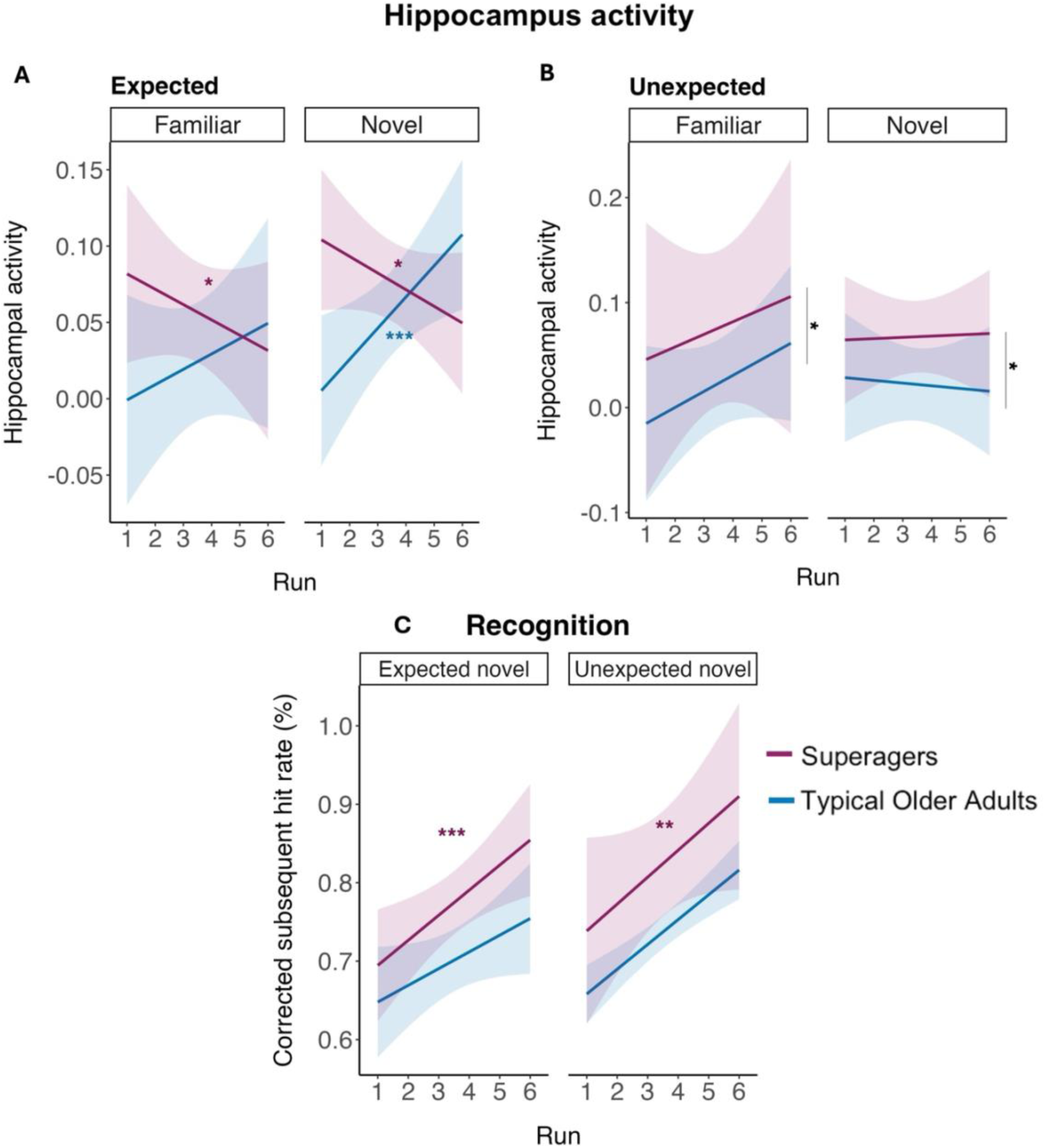
Encoding fMRI activity in the hippocampus and recognition performance. **A**, Hippocampal activity for expected events across experimental runs. Superagers show decreased hippocampal activation for expected events over time, regardless of their novelty status (familiar and novel), whereas typical older adults show increased hippocampal activation for expected events over time regardless of their novelty status (both for familiar and novel images). **B**, Hippocampal activity for unexpected events across experimental runs. Superagers show higher hippocampal activity than typical older adults for unexpected events, regardless of their novelty status (black asterisks, p = 0.01). **C**, Recognition performance for expected and unexpected images seen throughout the experimental runs. Superagers show better recognition performance for expected (p < 0.001) and unexpected images (p = 0.001) viewed in the last run compared to images viewed in the first run. Error bars represent 95% confidence intervals. Coloured asterisks denote significance levels for the simple slopes analysis. *p < 0.05, **p<0.01,***p < 0.001.

There is mounting evidence that neural responses in visual processing areas reflect a combination of repetition suppression arising from the encoding of stimulus probabilities and suppression effects driven from the encoding of mismatches between those predictions and their outcomes (Egner et al., 2010; Todorovic et al., 2012). To dissociate expectation from repetition suppression effects, we also tested expectation effects in the hippocampus independently of novelty while controlling for stimuli repetition (factors: group, expectation, and run + number of repetitions). This control analysis yielded the same pattern of results: superagers still exhibited decreased hippocampal activation for expected events over time (t = −2.19, p = 0.03), whereas typical older adults exhibited increased activation over time (t = 4.71, p < 0.001). These results suggest that expectation-related hippocampal suppression in superagers cannot be explained merely by low-level mechanisms associated with repeated presentations of the stimuli.

In turn, for unexpected events (Figure 3B), we found a main effect of group (β = – 0.024, χ²_(1)_ = 6.49, p = 0.01), with superagers showing greater hippocampal activation than typical older adults, consistently across runs. No other significant effects were found (all p’s > 0.11). These results suggest that for expected events, superagers show a pattern of hippocampal habituation that is independent of item novelty, consistent with expectation suppression, a pattern not observed in typical older adults. In contrast, for unexpected events, superagers exhibit greater hippocampal activation than typical older adults, also independent of item novelty. To assess whether these effects reflect an efficient encoding process in superagers, potentially leading to improvements in memory performance, we tested if group differences in subsequent recognition would emerge over time (experimental runs). To this end, we ran a mixed-effects model on subsequent recognition (factors: group, expectation, and run + recognition trial). This analysis revealed main effects of group (β = –0.25, χ²_(1)_ = 4.05, p = 0.044), expectation (β = –0.13, χ²_(1)_ = 10.41, p = 0.001), and run (β = –0.57, χ²_(5)_ = 60.17, p < 0.001) on memory performance. We also found an interaction between group and run (β = 0.15, χ²_(5)_ = 11.61, p = 0.040) (Figure 3C). No other interactions reached significance (all p’s > 0.25). Pairwise comparisons revealed that superagers had better recognition of items viewed in the last run compared to the first run for both expected (t = -5.01, p _FDR_ < 0.0001), and unexpected images (t = -4.21, p _FDR_ = 0.001), whereas in typical older adults, this superior performance was at trend level for expected (t = -2.37, p _FDR_ = 0.06) and unexpected images (t = -2.35, p _FDR_ = 0.07). Irrespective of expectation, group differences in recognition were most pronounced for images presented in the final run (t = -2.85, p _FDR_ = 0.004), with superagers outperforming typical older adults, whereas no group differences were observed for images viewed in the first run (t = - 0.68, p _FDR_ = 0.49), indicative of a stronger recency effect in superagers. Taken together, these findings suggest that superagers exhibit hippocampal habituation of expected information over exposure and a stronger hippocampal response to expectation violations, with superior subsequent recognition performance that is maintained throughout the encoding task.

#### Differential midbrain responses to contextual novelty in superagers vs. typical older adults

We next proceeded to explore expectation and novelty effects in the dopaminergic midbrain. For SN/VTA activation (Figure 4A), a 2 x 2 x 2 linear mixed effects model (factors: group, expectation and novelty) showed a marginal effect of expectation (β = 0.011, χ²_(1)_ = 3.40, p = 0.06) and novelty (β = -0.011, χ²_(1)_ = 3.37, p = 0.06). No significant main effect of group was found (β = -0.017, χ²_(1)_ = 1.03, p = 0.30). We found a significant interaction between group and novelty (β = -0.018, χ²_(1)_ = 8.66, p = 0.003) and a three-way interaction between group, expectation and novelty (β = 0.012, χ²_(1)_ = 3.99, p = 0.04). The rest of the interactions were not significant (all ps >0.40). Planned pairwise comparisons of the significant three-way interaction revealed that typical older adults exhibit decreased midbrain activation for unexpected familiar compared to unexpected novel objects (t = -2.67, p _FDR_ = 0.02), as well as a trend for decreased midbrain activation for unexpected compared to expected familiar objects (t = -2.23, p _FDR_ = 0.05). Typical older adults also showed decreased midbrain activation for familiar compared to novel objects regardless of their expectation (t = -3.32, p _FDR_ = 0.005). Compared to superagers, typical older adults showed decreased midbrain activity for unexpected familiar objects (t = - 2.42, p _FDR_ = 0.01). These findings suggest that typical older adults show a reduction in SN/VTA activity for unexpected familiar objects.

**Figure 4.**
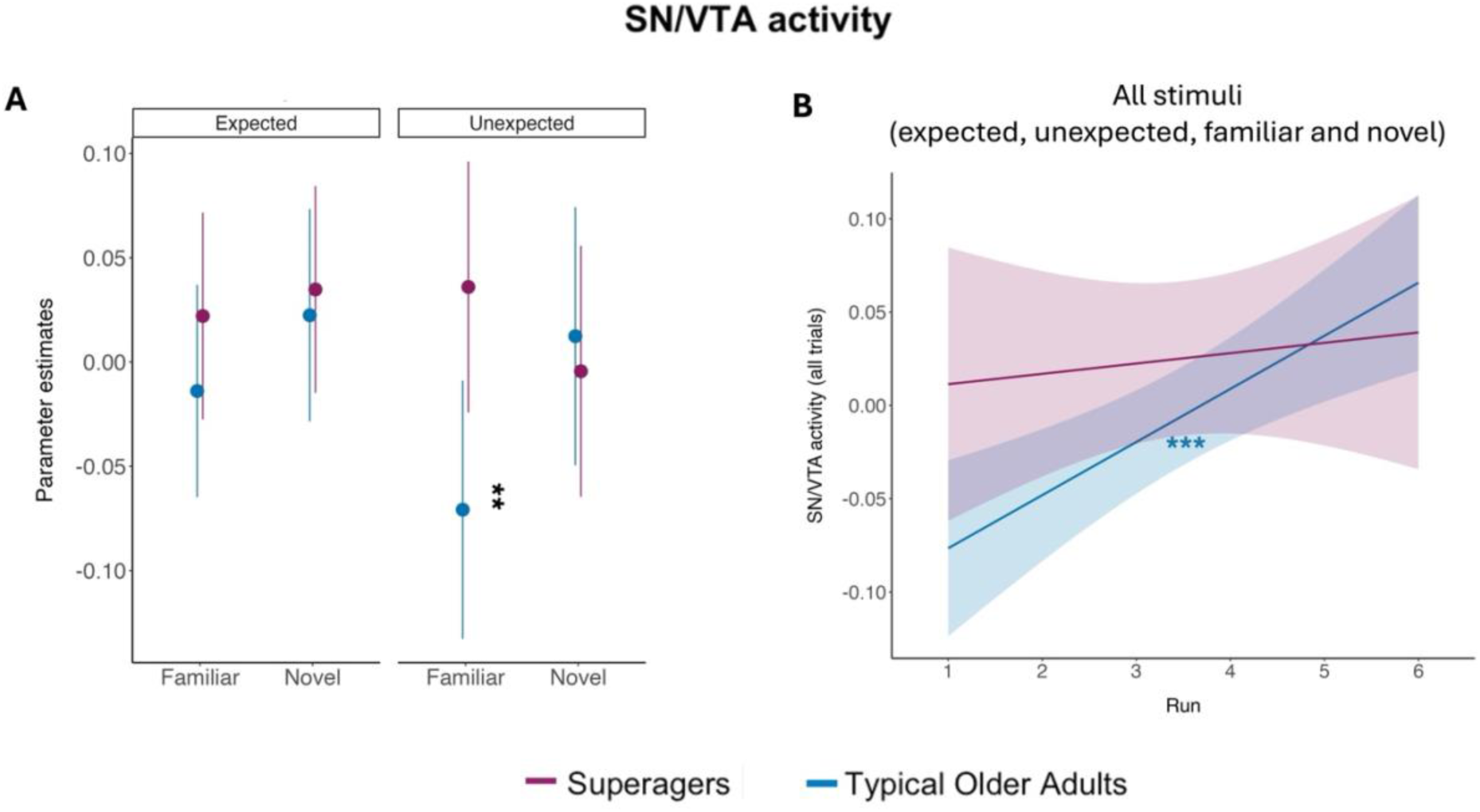
Encoding fMRI activity in the midbrain. **A**, Mean SN/VTA activity for each group (superagers vs. typical older adults) across the four experimental conditions defined by expectation (expected vs. unexpected) and item novelty (familiar vs. novel). Typical older adults exhibit decreased midbrain activation for unexpected familiar objects compared to expected novel objects (black asterisks, p = 0.006). **B**, Midbrain activity across experimental runs for all stimuli (expected, unexpected, familiar and novel). Typical older adults show a marked increase in SN/VTA activity over experimental runs (p < 0.001), with superagers having a stable response across the experiment (p = 0.19), with no significant expectation or novelty effects developing over time. Error bars represent 95% confidence intervals. Coloured asterisks denote significance levels for the simple slopes analysis. **p < 0.01, *** p < 0.001.

We next tested for repetition effects present in the midbrain, as these have been previously reported (Bunzeck et al., 2006, 2014; Zeithamova et al., 2016), with the use of a mixed-effects model (factors: group, expectation, novelty and run). We found a main effect of run (β = -0.05, χ²_(5)_ = 30.04, p < 0.001), with SN/VTA activity increasing across runs. Furthermore, we observed a significant group by run interaction (β = 0.009, χ²_(5)_ = 14.31, p = 0.014), driven by a marked increase in SN/VTA activity across runs in typical older adults (Figure 4B). Post hoc comparisons of SN/VTA slopes revealed a significant increase in SN/VTA activity over experimental runs only for typical older adults (t = 4.42, p _FDR_ < 0.001), with superagers having a stable response across the experiment (t = 1.32, p _FDR_ = 0.19). Typical older adults started with lower SN/VTA activity than superagers but gradually increased until reaching a level comparable to that of superagers by the end of the experiment. Replicating the findings from the first SN/VTA activity model, including run effects in the model did not change the significant interaction between group and novelty (β = -0.01, χ²_(1)_ = 8.77, p = 0.003), and the three-way interaction among group, expectation, and novelty was also significant (β = 0.01, χ²_(1)_ = 4.06, p = 0.044). Similarly, there were trends toward significance for the main effects of expectation (β = 0.01, χ²_(1)_ = 3.25, p = 0.071) and novelty (β = -0.01, χ²_(1)_ = 3.43, p = 0.064). The rest of the interactions were not significant (all ps >0.2). These findings suggest that SN/VTA activity is mainly modulated by group and the effects of novelty, with no significant habituation to expectation or novelty across time (i.e., no repetition suppression effects in the midbrain as a factor of novelty or expectation). Altogether, in response to expectation violations, typical older adults exhibit a novelty-driven processing in the hippocampus and midbrain.

#### Neuromelanin accumulation does not account for differences in midbrain activation

We continued by investigating whether the observed reduction in SN/VTA activity in response to unexpected familiar objects in typical older adults, compared to superagers, reflects impaired processing of contextual novelty (i.e., expectation violations). This reduction may be indicative of age-related decline in midbrain function, an effect that may also be evident in structural markers. To do so, we used NM accumulation as a proxy for dopaminergic system integrity, given that NM is a byproduct of dopamine metabolism. NM accumulation is assayed by CNR levels in the SN/VTA area, where higher CNR levels reflect higher NM accumulation. We conducted a linear regression model focusing on the relationship between each participant’s CNR and their SN/VTA BOLD responses to unexpected vs expected familiar items (i.e., the comparison showing the largest BOLD difference in Figure 4A), controlling for age and TIV [SN/VTA unexfam > SN/VTA expfam ∼ group * CNR + Age + TIV] (Figure 5A). The analysis revealed a main effect of CNR (*F*_(1,35)_ = 16.32, *p* = 0.0002, η² =0.32), however, neither the main effect of group (*F*_(1,35)_ = 1.20, *p* = 0.27, η² = 0.03) nor the interaction between group and CNR was significant (*F*_(1,35)_ = 0.13, *p* = 0.71, η² = 0.003). Follow-up analyses revealed that stronger unexpected familiar responses in SN/VTA were associated with lower CNR values both for superagers (t = -3.08, p _FDR_ < 0.001) and typical older adults (z = -2.64, p _FDR_ = 0.01). Surprisingly, these results indicate that reduced midbrain response to unexpected familiar items is associated with higher NM levels.

**Figure 5.**
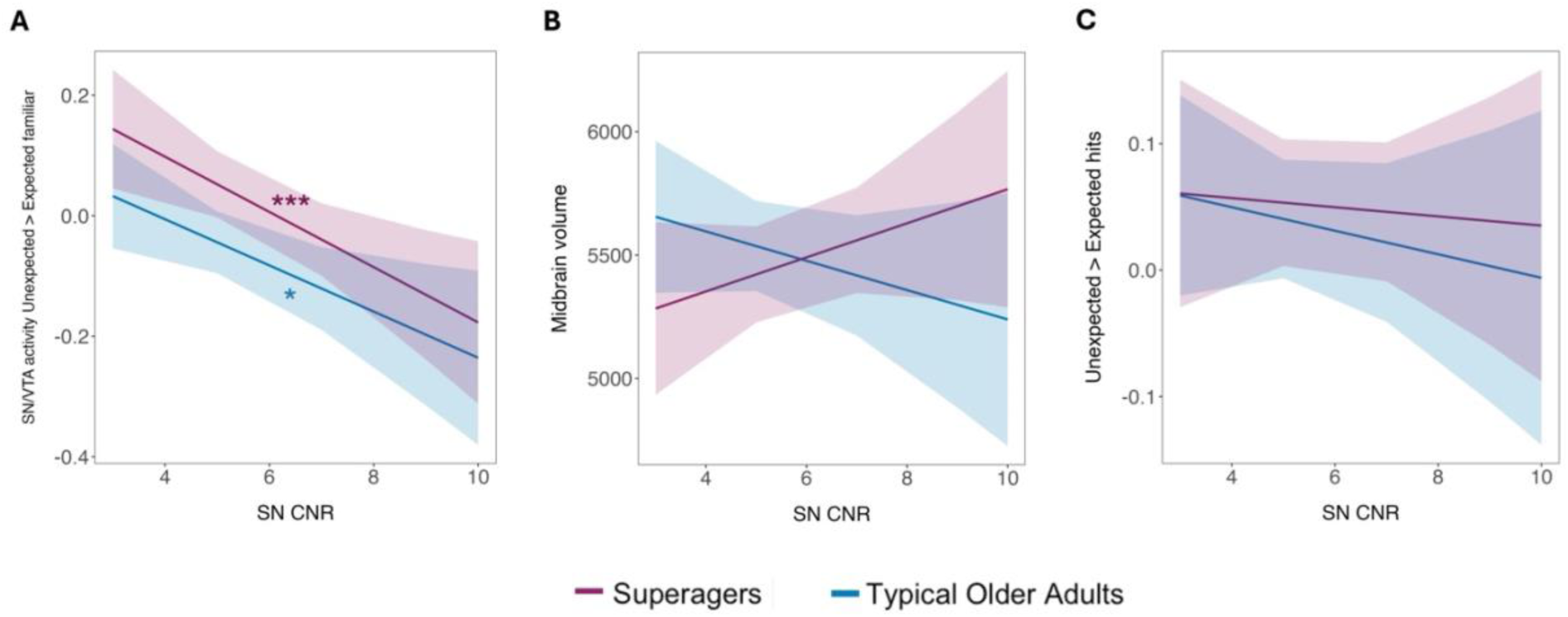
Neuromelanin interactions with midbrain activity, volume and memory performance on our task. **A**, In the SN/VTA the BOLD response to unexpected familiar objects is higher for individuals with lower levels of NM, both in typical older adults (p = 0.01) and in superagers (p < 0.010). **B**, Superagers and typical older adults show a trend towards and opposite relationship between neuromelanin accumulation and midbrain volume (p = 0.09). **C**, Relationship between NM accumulation with the unexpected behavioural outcome (subtracting the expected signal, with positive values reflecting a stronger unexpected signal and negative values reflecting a stronger expected signal). Suggesting that the effect of NM accumulation on memory outcomes did not differ between groups. Error bars represent 95% confidence intervals. Coloured asterisks denote significance levels for the simple slopes analysis.*p < 0.05, ***p<0.001

Considering NM accumulation as an index of dopaminergic system integrity, we expected it to be associated with other structural markers, such as bigger midbrain volume and better memory performance. However, we did not observe differences in SN/VTA volume between superager and typical older adults as a function of putative integrity of dopaminergic neurons. We implemented a linear regression model examining the relationship between NM accumulation and midbrain volume (Figure 5B), treating both measures as indices of dopaminergic integrity [Midbrain volume ∼ Group * CNR + Age +TIV]. There was a marginal main effect of group (*F*_(1,35)_ = 3.05, *p* = 0.08, η² < 0.08) and for the interaction between group and CNR (F_(1,35)_ = 3.03, p = 0.09, η² = 0.08). The main effect of CNR was not significant (*F*_(1,35)_ = 0.01, *p* = 0.89, η² < 0.001). These results reveal a trend that the relationship between NM accumulation and midbrain volume differs by group, with superagers showing a positive association consistent with preserved dopaminergic integrity. To examine whether NM accumulation was related to performance in our experimental memory task, we conducted a linear regression model examining the strength of the expectation violation memory boost observed behaviourally (subtracting expected hits from the unexpected ones) as a function of CNR values [unexpected hits > expected hits ∼ group * CNR + Age + TIV] (Figure 5C). However, we did not find significant effects of group (*F*_(1,35)_ = 0.01, *p* = 0.89, η² = 0.001), CNR (*F*_(1,35)_ = 0.46, *p* = 0.50, η² < 0.001), or the interaction between group and CNR (*F*_(1,35)_ = 0.08, *p* = 0.76, η² = 0.002). These findings indicate that memory performance for expectation violations was not directly associated with NM accumulation. Similarly, NM accumulation was not associated with general episodic memory performance, as the interaction between group and CNR on FCSRT scores was also not significant (Figure S2). Additional neuromelanin analysis, including the absence of group differences in baseline NM and age interactions (Figure S3, S4) as well as voxel-wise group comparisons (Figure S5) are included in the Supplemental Materials. Overall, our findings suggest that group differences in midbrain structure, activity, and memory performance are not related to neuromelanin accumulation.

## DISCUSSION

In an fMRI experiment, we manipulated item and contextual novelty at encoding to investigate whether superagers’ superior memory performance extended to contextual novelty and whether this would be supported by a hippocampal-midbrain encoding mechanism that resists typical ageing. Behaviourally, superagers showed overall better recognition performance, but both superagers and typical older adults exhibited enhanced recognition for unexpected compared to expected novel events, indicating preserved responses to expectation violations in both groups. At the brain structural level, we replicated previous findings (Harrison et al., 2018; Sun et al., 2016; Garo-Pascual et al., 2023) showing that superagers have preserved grey matter volumes in key memory-related regions of the medial temporal lobe. Functionally, during encoding superagers displayed increased hippocampal responses to contextual expectation violations consistently across runs compared to typical older adults. Interestingly, superagers showed hippocampal habituation to contextually expected stimuli across runs, irrespective of item novelty. Typical older adults, on the other hand, exhibited increasing hippocampal responses for expected novel items, as a function of run. In the midbrain, typical older adults showed reduced responses to unexpected familiar items compared to superagers. However, neuromelanin accumulation, as a proxy of dopaminergic integrity, did not modulate the group differences observed for midbrain activity and memory performance. Overall, these results suggest that superagers demonstrate an adaptive hippocampal function, that efficiently suppresses responses to expected information while remaining highly responsive to violations. Conversely, typical older adults exhibit reduced midbrain responses to unexpected familiar items, and an increased hippocampal response to expected novel items, suggesting a novelty-oriented processing.

Our behavioural results align with evidence that contextual expectation violations enhance recognition memory in young adults (Wittmann et al., 2007; Kafkas & Montaldi, 2015, 2018; Frank et al., 2020, 2022) and extend this effect to cognitively healthy older adults over the age of 80. Notably, although superagers demonstrated overall superior memory performance, extending observations from standardised neuropsychological assessment, both groups benefited from the expectation violation memory enhancement. This suggests that contextual novelty processing is preserved in healthy ageing and that group differences may be less pronounced than initially anticipated, or may rely on different processes to achieve the same recognition memory outcome (e.g., recollection vs. familiarity). Nevertheless, we did observe different patterns of neural engagement between groups during encoding. Across encoding runs, superagers showed stronger hippocampal responses to unexpected items compared to typical older adults. These results are in line with previous findings in young adults showing increased hippocampal responses to expectation violations that improve subsequent recognition (Wittmann et al., 2007; Kafkas & Montaldi, 2015, 2018; Frank et al., 2020, 2022).

Expectation effects typically reflect aggregated responses for mis/matched stimuli (as shown above); however, expectation can also unfold over time. In such scenarios, as the task progresses, expectation can engage predictive and habituation processes (Strange et al., 1999; Strange and Dolan, 2001; Yamaguchi et al., 2004; Murty et al., 2013). We found that superagers exhibited hippocampal habituation (reduced response) to expected events as the task progressed, reflecting an expectation suppression effect (Summerfield & de Lange, 2014). Superagers’ habituation to expectation could reflect heightened sensitivity to the cue-outcome contingency, reinforcing their correct prediction on expected trials (Summerfield et al., 2008), irrespective of item novelty. Previous studies have shown that expectation suppression effects can be dissociated from stimulus repetition effects in areas of the ventral visual system such as the posterior parahippocampal gyrus (Richter et al., 2018), and the fusiform face area (Egner et al., 2010; Larsson & Smith, 2012; Todorovic et al., 2012). In line with hierarchical predictive coding theories (Friston, 2005; Auksztulewicz and Friston, 2016), superagers may therefore exhibit a more efficient predictive mechanism in the hippocampus, characterised by adaptive neural suppression of expected events while remaining highly sensitive to unexpected stimuli (Richter et al., 2022). This mechanism may underlie superagers’ preserved memory abilities, however we could not relate these encoding effects to subsequent memory performance in our task (analysis reported in Supplemental Materials).

In typical older adults the pattern of results suggests item novelty played a stronger role than contextual novelty, as hippocampal responses to expected novel items increased as the task progressed. Furthermore, in the midbrain, typical older adults showed a reduced response to unexpected familiar items. This finding is somewhat surprising, as midbrain activity is often positively associated with expectation violation (Wittmann et al., 2007; Kafkas & Montaldi, 2015). However, when considered alongside the hippocampal effect, these findings suggest that typical older adults’ neural responses were more sensitive to item novelty than contextual novelty. In unexpected familiar trials, the cue was predicting a novel item, but was unexpectedly followed by a familiar item, therefore, the reduced midbrain response in typical older adults may reflect an omission of novelty response. In the reward domain, omission of a predicted reward suppresses dopamine neuron firing, constituting a negative prediction error (Schultz et al., 1997; Hollerman & Schultz, 1998). Given that the SN/VTA responds to novelty in ways that parallel reward processing (Bunzeck & Düzel, 2006; Wittmann et al., 2007), and that novelty has been proposed to function as a motivational signal within the hippocampal-VTA loop (Lisman & Grace, 2005; Kakade & Dayan, 2002), the absence of expected novelty on these trials may have produced an analogous suppression. This account would suggest that typical older adults still track the cue prediction, but allocate their resources more towards novel information. An alternative explanation is that the attenuated midbrain response in typical older adults reflects a more general decrease in dopaminergic efficiency rather than a novelty-specific omission signal. Age-related reductions in dopamine synthesis, receptor density, and transporter availability are well-documented (Bäckman et al., 2006; Li et al., 2010) and affect the midbrain-hippocampal circuitry that supports novelty detection and memory encoding (Düzel et al., 2008; Chowdhury et al., 2012).

To examine these possibilities, we used neuromelanin-sensitive MRI as a proxy for functional integrity of the dopaminergic system in the brain (Zucca et al., 2017). Although we did not find any group-specific relationships between neuromelanin accumulation and midbrain activation or memory performance, we observed a trend towards opposite group trajectories relative to midbrain volume. This may suggest potentially distinct neuromelanin profiles in ageing, consistent with animal model findings that indicate a dual role for neuromelanin; while it provides intracellular neuroprotection, it may reflect neurotoxicity when released extracellularly following neuronal death (Zecca et al., 1996; Zucca et al., 2017, 2018). Future studies accounting for individual baseline differences and longitudinal designs would be needed to clarify the role of neuromelanin in age-related dopaminergic function. While our results are generally in line with previous contextual expectation effects on memory, and extend them as expectation unfolds in time, the item novelty effects observed were less pronounced. Previous findings in young adults found anticipatory hippocampal and midbrain responses linked to enhanced memory formation (Wittmann et al., 2007; Poh et al., 2022). However, our experiment was not designed to dissociate anticipatory from outcome-related mechanisms. To implicitly induce expectation violations, the cue and outcome were presented in close temporal proximity and orthogonally to the encoding task instructions (manmade/natural decision). Consequently, our encoding results may reflect a combined contribution of anticipatory and novelty-related hippocampal and midbrain responses. Alternatively, additional neuromodulatory influences known to support memory for novel information in younger adults, such as noradrenergic and cholinergic inputs (Ranganath & Rainer, 2003; Meeter, Murre, & Talamini, 2004; Hasselmo and Stern, 2006; Kafkas & Montaldi, 2018b; Kafkas, 2021) that could also be contributing to our results. In line with this possibility, recent evidence suggests that superagers maintain preserved cholinergic innervation, as they exhibit significantly fewer neurofibrillary tangles in the basal forebrain cholinergic system (Weintraub et al., 2025), which may contribute to the preservation of memory function in late life. Future work could examine the effects of such neuromodulatory influences on cognitive functioning, to better understand the mechanisms supporting superior memory performance in superagers.

In conclusion, we provide novel evidence that superagers and typical older adults engage hippocampal-midbrain circuitry differently when processing expected and unexpected information during encoding. Superager exhibited heightened hippocampal sensitivity to expected and unexpected information. This pattern is consistent with an efficient predictive mechanism that suppresses redundant input while remaining responsive to expectation violations. Typical older adults showed a different, more novelty-oriented pattern, with hippocampal engagement selectively increasing for anticipated novel content and attenuated midbrain response to absence of novelty. Although the direct link between these encoding dynamics and subsequent recognition performance remains to be established, the present findings identify hippocampal prediction processing as a promising candidate mechanism for understanding what distinguishes exceptional from typical cognitive ageing, and motivate future work examining whether interventions that support predictive efficiency could benefit memory function in later life.

## Acknowledgments

We thank the participants of the Vallecas Project and the staff of the CIEN Foundation. We thank Marta Garo-Pascual for helpful discussions and valuable feedback on earlier versions of this manuscript. This work was supported by the CIEN Foundation and the Queen Sofia Foundation, as well as a grant from the Spanish Ministry of Science and Innovation to B.S. (PID2023-153259OB-I00). This work was supported by a Marie Skłodowska-Curie fellowship to D.F. project PCI2021-122046-2B, financed by the Spanish Ministry of Science and Innovation and the Spanish State Research Agency MCIN/AEI/10.13039/501100011033 and the European Union “NextGenerationEU”/PRTR. D.F. is supported by a Royal Society University Research Fellowship URF/R1/241499. MGH is supported by Grant PRE2021-101023 funded by MCIN/AEI/10.13039/501100011033 and by ESF+. For the purpose of Open Access, the author has applied a CC BY public copyright license to any Author Accepted Manuscript version arising from this submission.

## Author contributions

D.F. and B.S. designed research; M.G.-H. and D.F. performed research; M.G.-H., L.Z-and D.F. analysed data; M.G.-H., L.Z., B.S. and D.F. wrote the paper.

## Data availability

Anonymised data can be accessed upon request to.

## Supplementary materials

### Rule-learning

We found significant differences in the number of sessions required to learn the rule (t_(39)_ = 3.4, p = 0.002, Cohen’s d = 1.06). Superagers required fewer sessions to learn the rule (mean = 1 session, median = 1 session, SD = 0.2) compared to typical older adults (mean = 1.7 sessions, median = 1.5 sessions, SD = 0.8).

### Encoding

Two subjects were not included in the analyses as they did not make any responses inside the scanner. A mixed 2 x 2 x 2 ANOVA (factors: group, expectation and novelty) for the manmade/natural decision accuracy in the encoding phase showed an interaction between expectation and novelty (F_(1,37)_ = 5.59, p = 0.02, η² < 0.001) and no main effect of group (F_(1,37)_ = 0.27, p = 0.60, η² = 0.007). Pairwise comparisons revealed that in the unexpected condition, accuracy was significantly higher for familiar compared to novel items (z = 2.25, p _FDR_ = 0.03) and no significant difference between familiar and novel conditions in the expected condition (z = 0.23, p _FDR_ = 0.81). There was no significant difference between expected and unexpected conditions when objects were familiar (z = -0.53, p _FDR_= 0.598). However, when objects were novel, accuracy was significantly higher for expected compared to unexpected condition (z = 2.65, p _FDR_ = 0.01). No other significant effects were found (al ps > 0.06). The effect was mainly driven by confusing stimuli (e.g., a manmade figure of a saint). Removing images that no participant categorised correctly eliminated the interaction between expectation and novelty (F(1,37) = 2.80, p = 0.10, η² < 0.001), and all other effects were nonsignificant (all ps > 0.29).

For reaction times in the encoding phase a repeated measures 2 x 2 x 2 ANOVA (factors: group, expectation and novelty) showed a main effect of novelty, with faster responses for novel objects (F_(1,37)_ = 60.37, p < 0.001, η² = 0.23), and a main effect of group (F_(1,37)_ = 8.89, p =0.005, η² = 0.12), with typical older adults responding faster than superagers. No other significant effects were found (all ps > 0.14). These results indicate that responses to novel images were faster compared to familiar images, and that, compared to typical older adults, superagers had a longer encoding period of both familiar and novel objects. Furthermore, we observed a trend (t(39) = −1.75, p = 0.089, Cohen’s d = −0.55) suggesting that a greater proportion of superagers (61%) noticed the manipulation inside the scanner (i.e., mismatched -unexpected- events) compared with typical older adults (35%).

**Table S1.** Behavioural data for the encoding phase (mean reaction times, RT, with SE), separately for controls and superagers.

|  | **Novel**<br>**expected** | **Novel**<br>**unexpected** | **Familiar**<br>**expected** | **Familiar**<br>**unexpected** |
| --- | --- | --- | --- | --- |
| **Superagers (n=21)** |  |  |  |  |
| Decision accuracy in % (SE) | 81.16 (0.009) | 79.24 (0.01) | 81.63 (0.009) | 81.81 (0.01) |
| RT in ms (SE) | 1430 (0.01) | 1440 (0.02) | 1600 (0.01) | 1620 (0.02) |
| **Controls (n=18)** |  |  |  |  |
| Decision accuracy in % (SE) | 74.24 (0.01) | 73.13 (0.02) | 74.12 (0.01) | 74.43 (0.01) |
| RT in ms (SE) | 1270 (0.01) | 1340 (0.03) | 1520 (0.02) | 1480 (0.02) |

### Whole-brain analysis

We also ran an exploratory voxel-wise univariate whole-brain analysis. First-level GLMs were computed for each participant, an event-related design matrix was created containing the five conditions of interest: cue, expected familiar outcome, unexpected familiar outcome, expected novel outcome, and unexpected novel outcome. Additional regressors were included to control for physiological noise using CompCor correction.

**Figure S1.**
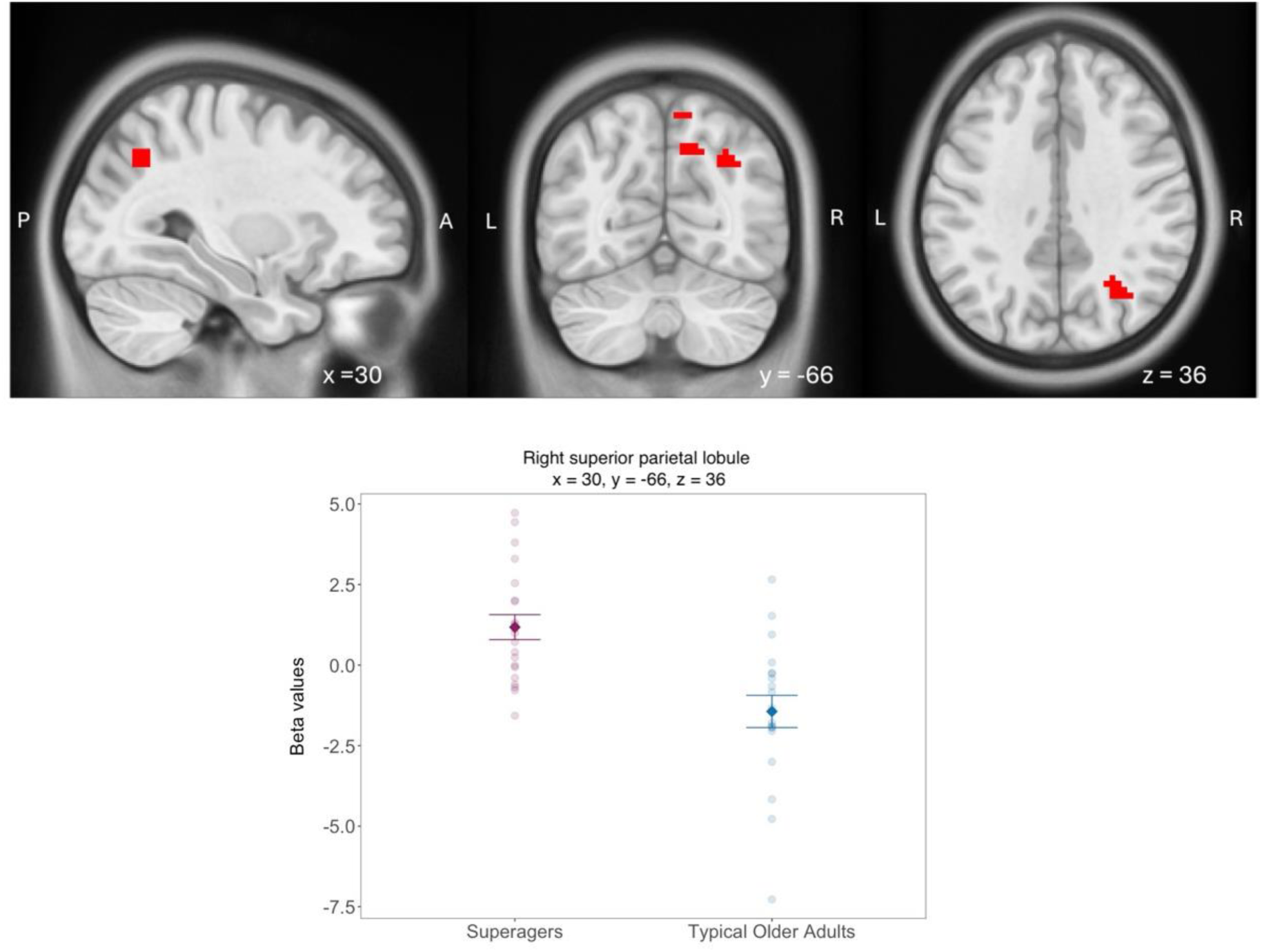
fMRI whole-brain group differences when processing expectation *violations overlaid in MNI space. Significant group differences of unexpected > expected familiar objects. Compared to typical older adults, superagers showed stronger activation in the right superior parietal lobule for unexpected compared to expected familiar objects. (*TFCE FWE-corrected p <0.05). Section coordinates refer to standard MNI space and are given in mm. A=anterior. P=posterior. L=left. R=right.

### Subsequent recognition

A subsequent memory analysis was performed using a 2 x 2 x 2 linear mixed-effects model (factors: group, expectation, and hits) to investigate whether encoding-related activity in the hippocampus and midbrain ROIs was associated with subsequent recognition performance. We did not find group differences in hippocampal (β = – 0.004, χ²(1) = 0.34, p = 0.55) or midbrain activity (β = –0.01, χ²(1) = 2.31, p = 0.12) at encoding that were associated with better subsequent recognition performance.

### Psychophysiological Interactions (PPI): Effective connectivity analysis

A PPI analysis with the SN/VTA and hippocampus masks as seed areas for unexpected versus expected familiar objects did not reveal significant group differences in connectivity.

### Neuromelanin analysis

To examine if preserved dopaminergic integrity is associated with better overall memory function, we examined its relationship with performance on the episodic memory test used to identify superagers (FSCRT; Figure S4). A linear regression model (FCSRT ∼ group * CNR + Age + TIV) showed a main effect of group (F_(1,35)_ = 7.22, p = 0.01, η² = 0.17). The main effect of CNR on FCSRT scores was not significant (F_(1,35)_ = 0.00, p = 0.99, η² < 0.001), and no interaction was found between group and CNR (F_(1,35)_ = 0.29, p = 0.58, η² = 0.008).

**Figure S2.**
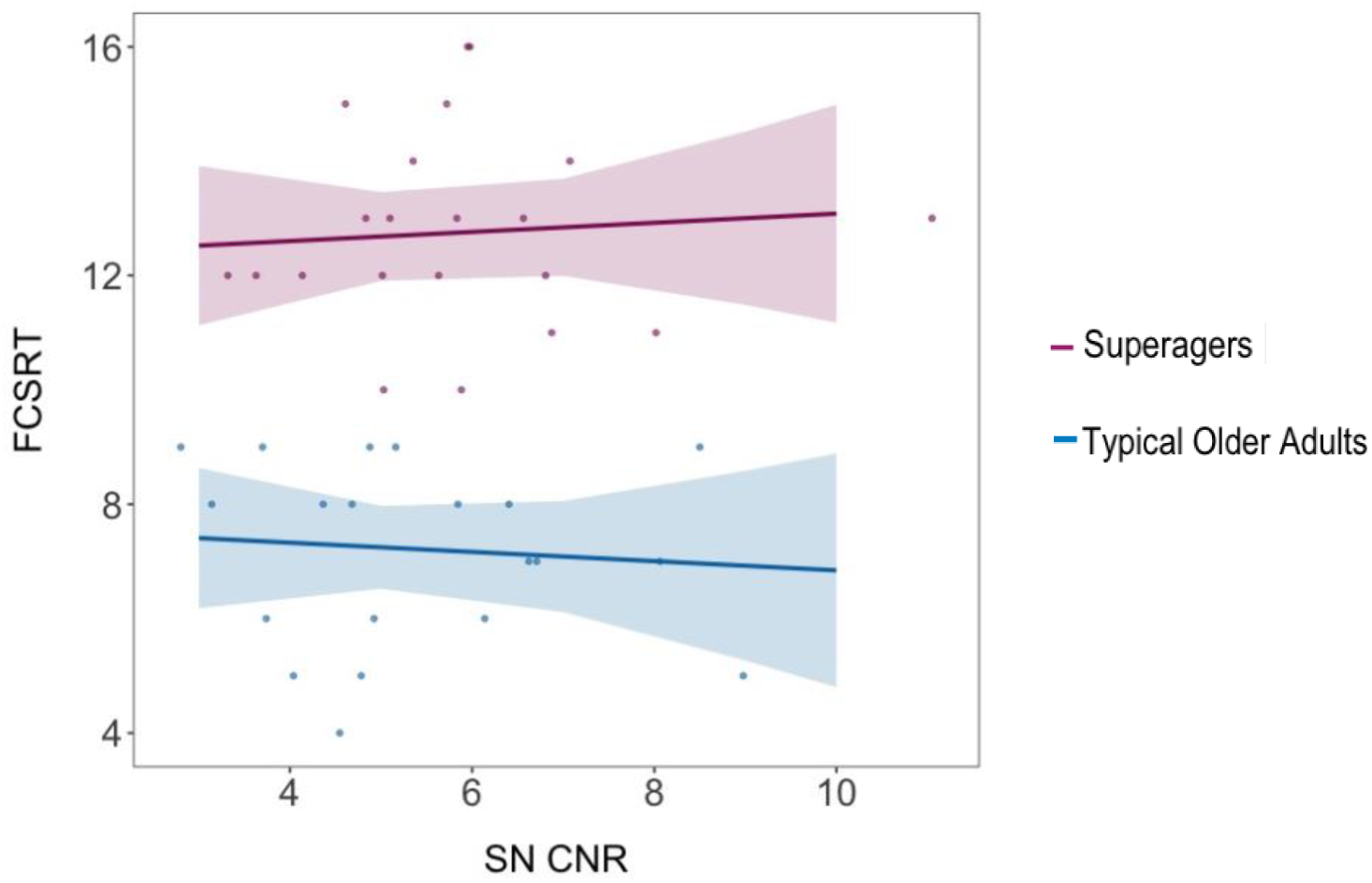
Neuromelanin accumulation and episodic memory performance (FCSRT test scores).

**Figure S3.**
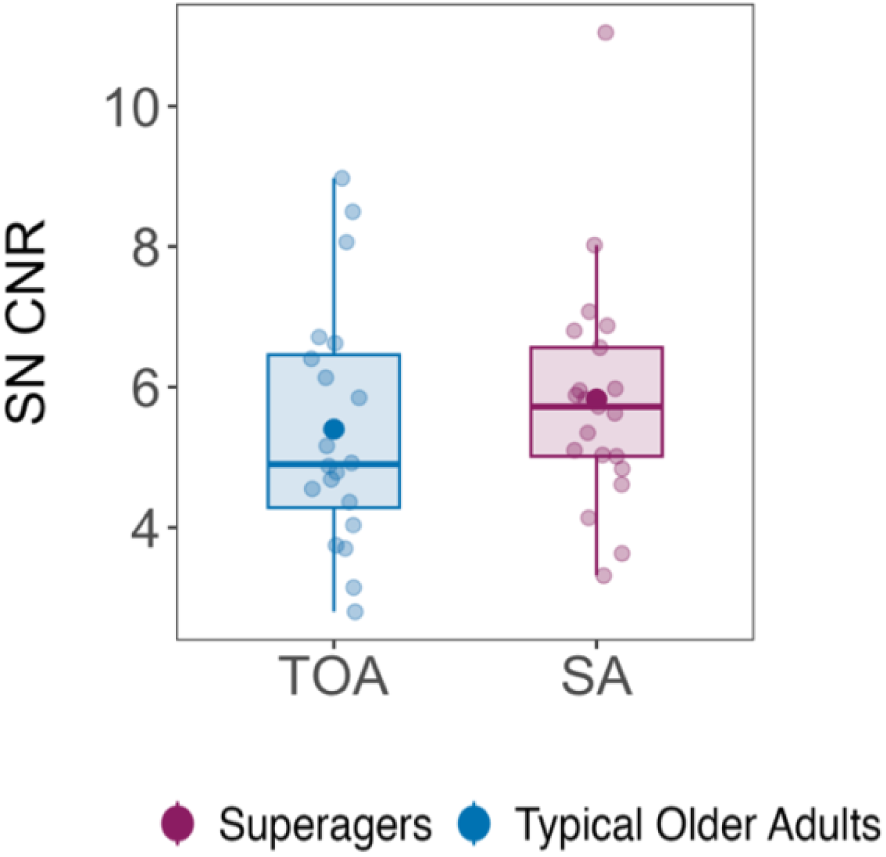
Neuromelanin accumulation across groups (boxplots show median ± IQR; mean indicated by a dot). We did not find significant group differences (t = 0.80, p = 0.42).

To examine age-related group trajectories in the preservation of midbrain volume, we tested group differences as a function of age. A linear regression model (*midbrain volume ∼ group * age + TIV*) revealed a main effect of age (*F*_(1,36)_ = 7.33, *p* = 0.01, η² = 0.17). Although superagers appeared to show a more stable midbrain volume across age, neither the main effect of group (*F*_(1,36)_ = 2.41, *p* = 0.12, η² = 0.06) nor the group and age interaction (*F*_(1,36)_ = 2.44, *p* = 0.12, η² = 0.06) reached statistical significance, likely due to the large variability observed.

**Figure S4.**
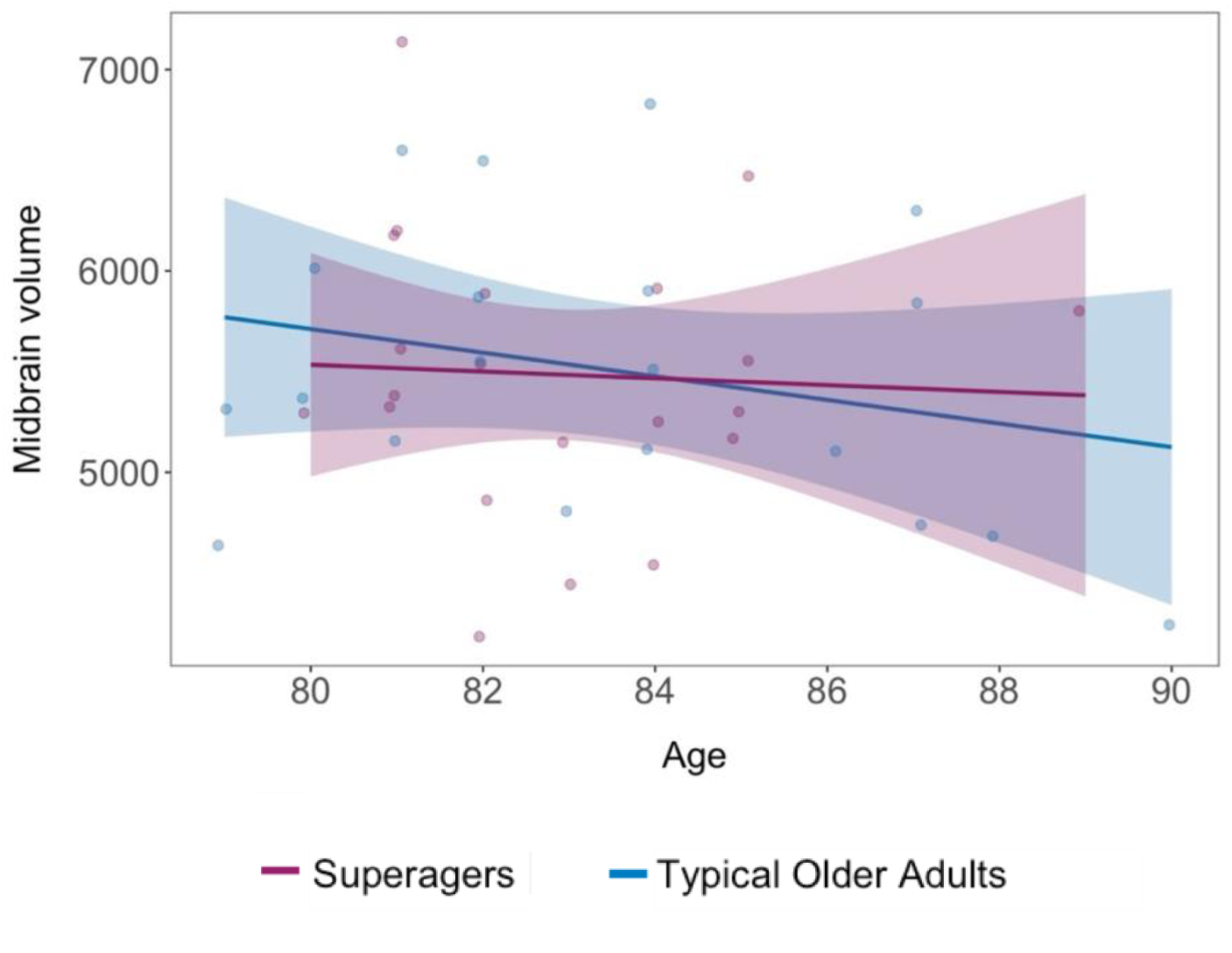
Midbrain volume as a function of age in superagers and typical older adults. Although superagers appeared to show a more stable trajectory across age, no significant group differences were observed.

**Figure S5.**
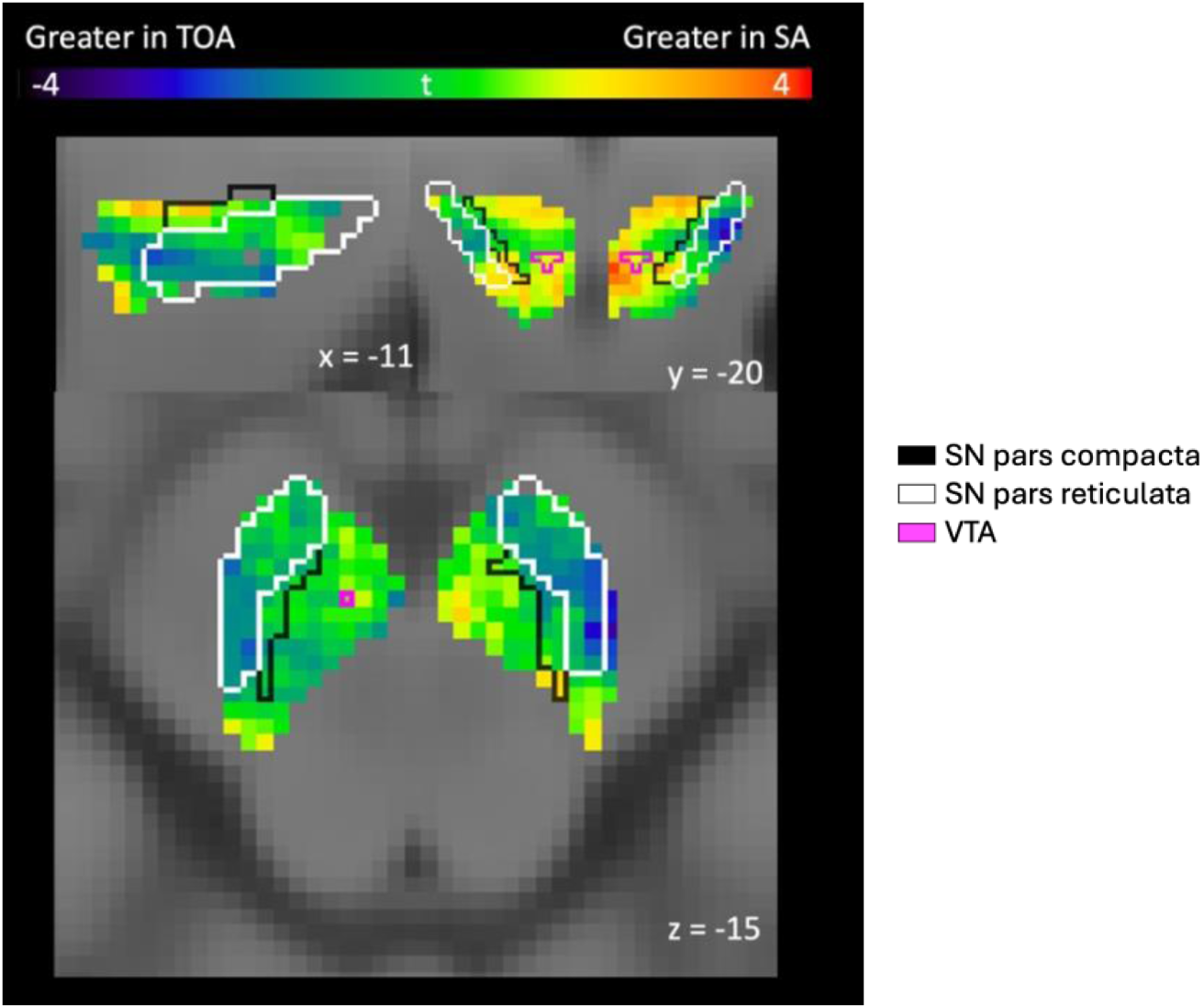
NM-MRI voxel-wise results. T-statistic map overlaid on the NM-MRI template illustrating SN CNR signal intensity in superagers vs typical older adults (permutation test, voxel level thresholded at p < 0.05, uncorrected). Warm colours (red/yellow) indicate higher t-statistic values and greater NM-MRI signal in superagers, whereas cool colours (blue) indicate higher signal in typical older adults. Superagers exhibit increased NM signal intensity in the medial substantia nigra pars compacta and regions adjacent to ventral tegmental area (VTA), while typical older adults show higher NM signal intensity in regions adjacent to the substantia nigra.

